# Glycosaminoglycans induce conformational change in the SARS-CoV-2 Spike S1 Receptor Binding Domain

**DOI:** 10.1101/2020.04.29.068767

**Authors:** Courtney J. Mycroft-West, Dunhao Su, Yong Li, Scott E. Guimond, Timothy R. Rudd, Stefano Elli, Gavin Miller, Quentin M. Nunes, Patricia Procter, Antonella Bisio, Nicholas R. Forsyth, Jeremy E. Turnbull, Marco Guerrini, David G. Fernig, Edwin A. Yates, Marcelo A. Lima, Mark A. Skidmore

## Abstract

The glycosaminoglycan (GAG) class of polysaccharides are utilised by a plethora of microbial pathogens as receptors for adherence and invasion. The GAG heparin prevents infection by a range of viruses when added exogenously, including the S-associated coronavirus strain HSR1 and more recently we have demonstrated that heparin can block cellular invasion by SARS-CoV-2. Heparin has found widespread clinical use as anticoagulant drug and this molecule is routinely used as a proxy for the GAG, heparan sulphate (HS), a structural analogue located on the cell surface, which is a known receptor for viral invasion. Previous work has demonstrated that unfractionated heparin and low molecular weight heparins binds to the Spike (S1) protein receptor binding domain, inducing distinct conformational change and we have further explored the structural features of heparin with regard to these interactions. In this article, previous research is expanded to now include a broader range of GAG family members, including heparan sulphate. This research demonstrates that GAGs, other than those of heparin (or its derivatives), can also interact with the SARS-CoV-2 Spike S1 receptor binding domain and induce distinct conformational changes within this region. These findings pave the way for future research into next-generation, tailor-made, GAG-based antiviral agents, against SARS-CoV-2 and other members of the *Coronaviridae*.

## Introduction

The glycosaminoglycans (GAGs) are polydisperse, heterogeneous natural products that are exploited as pharmaceuticals and nutraceuticals. Clinically, unfractionated heparin (UFH), low molecular weight heparins (LMWHs) and heparinoids are approved as anticoagulants/thrombotics with excellent safety, stability, bioavailability and pharmacokinetic profiles. Crucially, GAGs and their derivatives, some of which lacking significant anticoagulant activity^1^, are an under-exploited antiviral drug class, despite possessing broad-spectrum activity against a multitude of distinct viruses, including *coronaviridae* and SARS-associated coronaviruses^2,3^, in addition to flaviviruses^4,5^, herpes^6^, influenza^7^ and HIV^8,9^.

Traditional drug development processes are slow and ineffective against emerging public health threats such as the current SARS-CoV-2 coronavirus outbreak which makes the repurposing of existing drugs a timely and attractive alternative. Heparin, a well-tolerated anticoagulant pharmaceutical, has been used safely in medicine for over 80 years and alongside its anticoagulant activities, this biomolecule possesses the ability to prevent viral infection, including from members of the *coronaviridae*^1^. Furthermore, the closely related glycosaminoglycan (GAG) member, heparan sulfate (HS), is known to bind CoV surface proteins and to be used by coronavirus for its attachment to target cells^10^.

Currently, there is a dearth of commercially available medicinal products designed to treat and/or prevent infections associated with the new SARS-CoV-2 coronavirus outbreak. Here, we describe preliminary observations for the ability of the SARS-CoV-2 S1 RBD to interact with a broad range of GAGs, in addition to those previously demonstrated for UFH and LMWH^2^.

## Methods & Materials

### 2.1 Recombinant expression of SARS-CoV-2 S1 RBD

Residues 330−583 of the SARS-CoV-2 Spike Protein (GenBank: MN908947) were cloned upstream of a N-terminal 6XHisTag in the pRSETA expression vector and transformed into SHuffle® T7 Express Competent *E. coli* (NEB, UK). Protein expression was carried out in MagicMedia™ *E. coli* Expression Media (Invitrogen, UK) at 30°C for 24 hrs, 250 rpm. The bacterial pellet was suspended in 5 mL lysis buffer (BugBuster Protein Extraction Reagent, Merck Millipore, UK; containing DNAse) and incubated at room temperature for 30 mins. Protein was purified from inclusion bodies using IMAC chromatography under denaturing conditions. On-column protein refolding was performed by applying a gradient with decreasing concentrations of the denaturing agent (6 - 0 M Urea). After extensive washing, protein was eluted using 20 mM NaH2PO4, pH 8.0, 300 mM NaCl, 500 mM imidazole. Fractions were pooled and buffer-exchanged to phosphate-buffered saline (PBS; 140 mM NaCl, 5 mM NaH2PO4, 5 mM Na2HPO4, pH 7.4; Lonza, UK) using a Sephadex G-25 column (GE Healthcare, UK). Recombinant protein was stored at −20°C until required.

### 2.2 Secondary structure determination of SARS-CoV-2 S1 RBD by circular dichroism spectroscopy

The circular dichroism (CD) spectrum of the SARS-CoV-2 S1 RBD in PBS was recorded using a J-1500 Jasco CD spectrometer (Jasco, UK), Spectral Manager II software (JASCO, UK) and a 0.2 mm pathlength, quartz cuvette (Hellma, USA) scanning at 100 nm.min^−1^ with 1 nm resolution throughout the range λ = 190 - 260 nm. All spectra obtained were the mean of five independent scans, following instrument calibration with camphorsulfonic acid. SARS-CoV-2 S1 RBD was buffer-exchanged (prior to spectral analysis) using a 5 kDa Vivaspin centrifugal filter (Sartorius, Germany) at 12,000 g, thrice and CD spectra were collected using 21 μl of a 0.6 mg.ml^−1^ solution in PBS, pH 7.4. Spectra of glycosaminoglycans were collected in the same buffer at approximately comparable concentrations, since these are disperse materials. Collected data were analysed with Spectral Manager II software prior to processing with GraphPad Prism 7, using second order polynomial smoothing through 21 neighbours. Secondary structural prediction was calculated using the BeStSel analysis server^11^.To ensure that the CD spectral change of SARS-CoV-2 S1 RBD in the presence of GAGs did not arise from simply from the addition of the GAG spectra (this class of carbohydrates are known to possess CD spectrum at high concentrations^12,13^), difference spectra were analysed for each GAG in order to verify that the change in the CD spectrum arose from a conformational change following binding to the test GAG.

## Results

### 3.1 Secondary structure determination of SARS-CoV-2 S1 RBD protein by circular dichroism spectroscopy

Circular dichroism (CD) spectroscopy detects changes in protein secondary structure that occur in solution using UV radiation. Upon binding, conformational changes are detected and quantified using spectral deconvolution^14^. Indeed, SARS-CoV-2 S1 RBD underwent conformational change in the presence of GAGs (Figures 1 - 6). These observed changes further demonstrate that the SARS-CoV-2 S1 RBD interacts with not only UFH and LMWHs, but also other members of the GAG family of carbohydrates.

**Figure 1:**
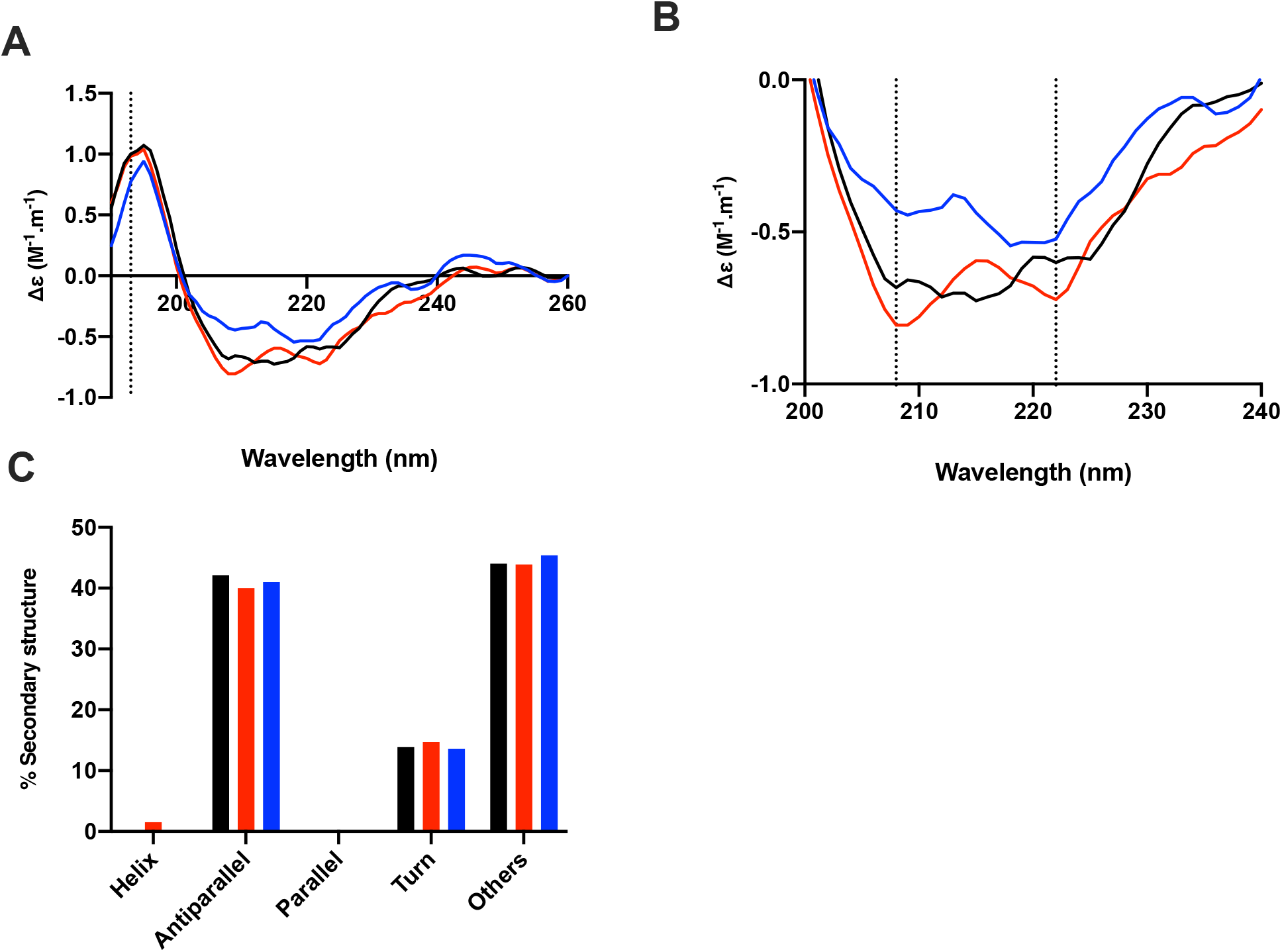
The conformational change of the SARS-CoV-2 S1 RBD observed in the presence of heparan sulphate by circular dichroism (CD) spectroscopy. (**A**) Circular dichroism spectra (190 - 260 nm) of nCovS1RBD alone (black solid line), heparin (red solid line) and heparan sulphate (blue) in PBS, pH 7.4. The dotted vertical line indicates 193 nm. (**B**) Details of the same spectra expanded between 200 and 240 nm. Vertical dotted lines indicate 222 nm and 208 nm. (**C**) Secondary structure content analysed using BeStSel for nCovS1RBD. α helical secondary structure is characterized by a positive band at ~193 nm and two negative bands at ~208 and ~222 nm (analysis using BeStSel was performed on smoothed data between 190 and 260 nm.

**Figure 2:**
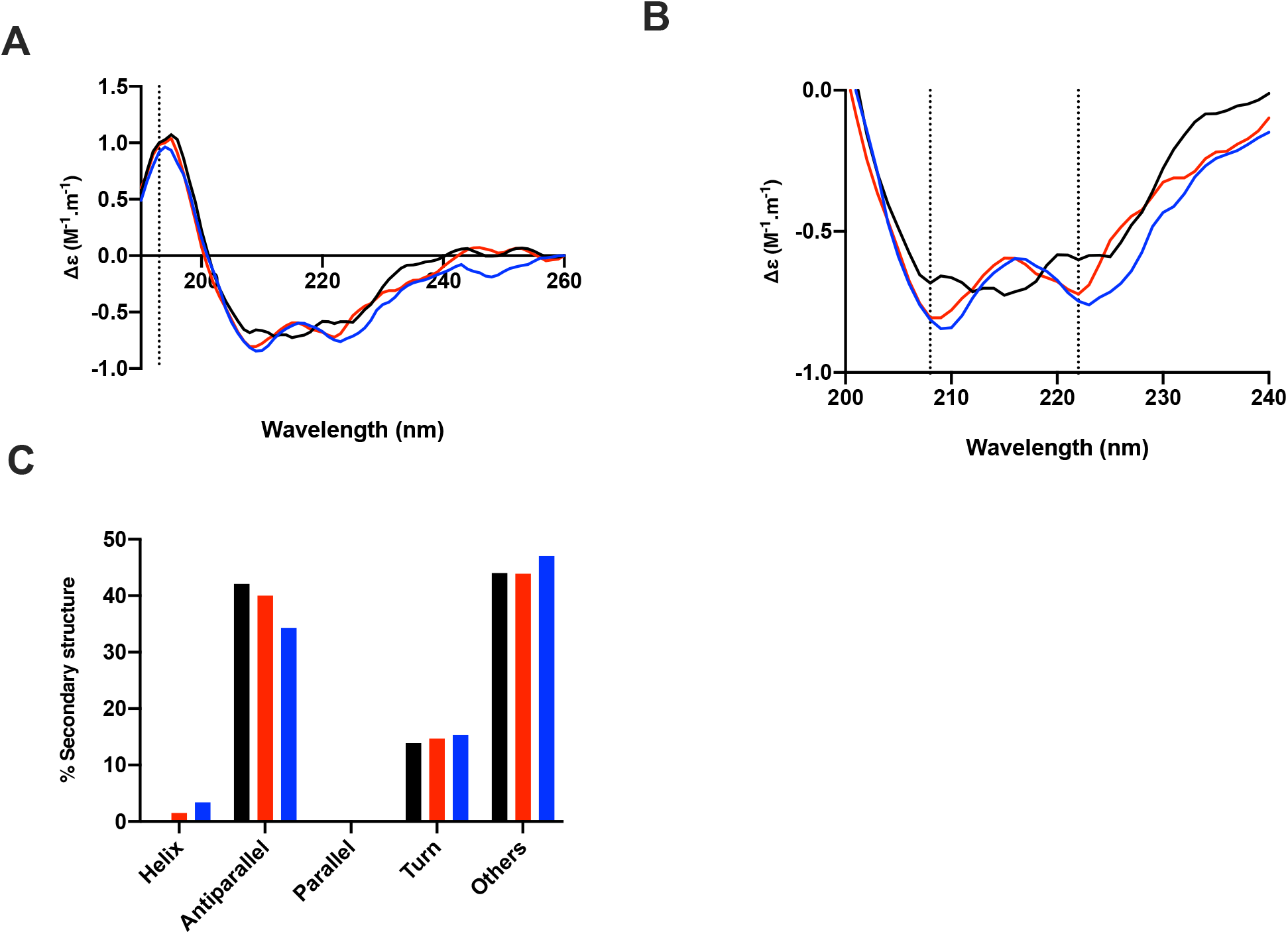
The conformational change of the SARS-CoV-2 S1 RBD observed in the presence of dermatan sulphate by circular dichroism (CD) spectroscopy. (**A**) Circular dichroism spectra (190 - 260 nm) of nCovS1RBD alone (black solid line), heparin (red solid line) and dermatan sulphate (blue) in PBS, pH 7.4. The dotted vertical line indicates 193 nm. (**B**) Details of the same spectra expanded between 200 and 240 nm. Vertical dotted lines indicate 222 nm and 208 nm. (**C**) Secondary structure content analysed using BeStSel for nCovS1RBD. α helical secondary structure is characterized by a positive band at ~193 nm and two negative bands at ~208 and ~222 nm (analysis using BeStSel was performed on smoothed data between 190 and 260 nm.

**Figure 3:**
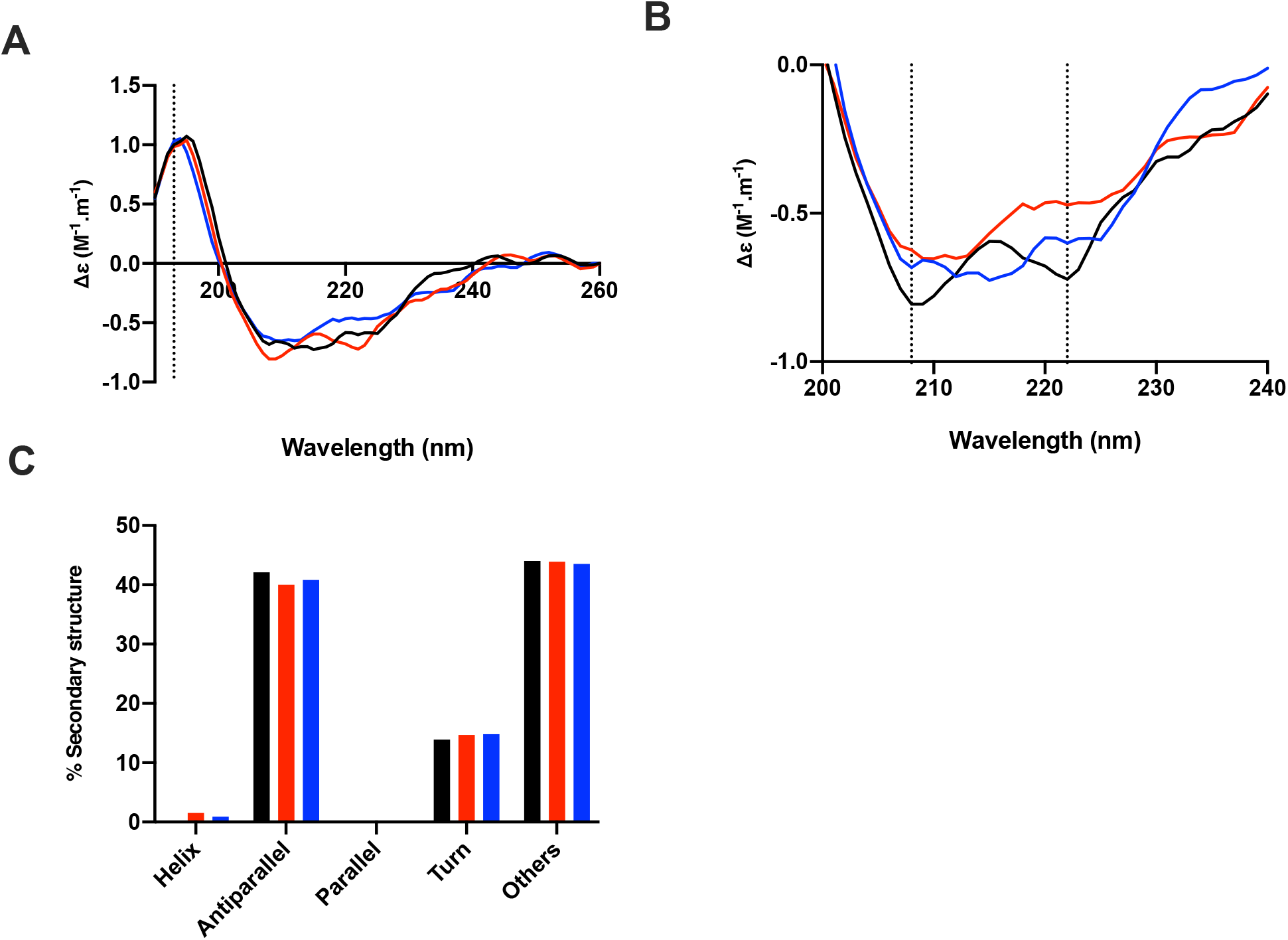
The conformational change of the SARS-CoV-2 S1 RBD observed in the presence of chondroitin sulphate A by circular dichroism (CD) spectroscopy. (**A**) Circular dichroism spectra (190 - 260 nm) of nCovS1RBD alone (black solid line), heparin (red solid line) and chondroitin sulphate A (blue) in PBS, pH 7.4. The dotted vertical line indicates 193 nm. (**B**) Details of the same spectra expanded between 200 and 240 nm. Vertical dotted lines indicate 222 nm and 208 nm. (**C**) Secondary structure content analysed using BeStSel for nCovS1RBD. α helical secondary structure is characterized by a positive band at ~193 nm and two negative bands at ~208 and ~222 nm (analysis using BeStSel was performed on smoothed data between 190 and 260 nm.

**Figure 4:**
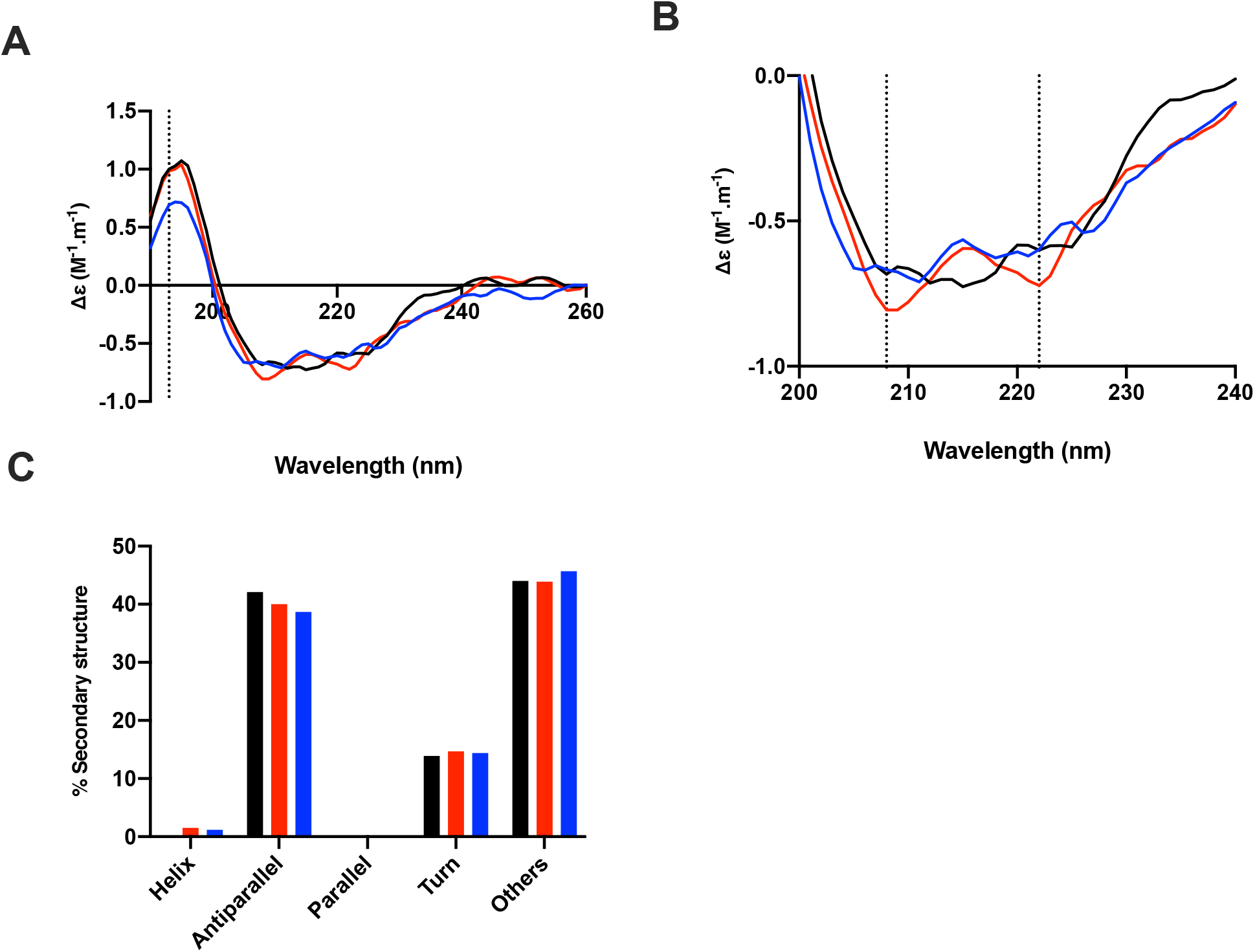
The conformational change of the SARS-CoV-2 S1 RBD observed in the presence of chondroitin sulphate C by circular dichroism (CD) spectroscopy. (**A**) Circular dichroism spectra (190 - 260 nm) of nCovS1RBD alone (black solid line), heparin (red solid line) and chondroitin sulphate C (blue) in PBS, pH 7.4. The dotted vertical line indicates 193 nm. (**B**) Details of the same spectra expanded between 200 and 240 nm. Vertical dotted lines indicate 222 nm and 208 nm. (**C**) Secondary structure content analysed using BeStSel for nCovS1RBD. α helical secondary structure is characterized by a positive band at ~193 nm and two negative bands at ~208 and ~222 nm (analysis using BeStSel was performed on smoothed data between 190 and 260 nm.

**Figure 5:**
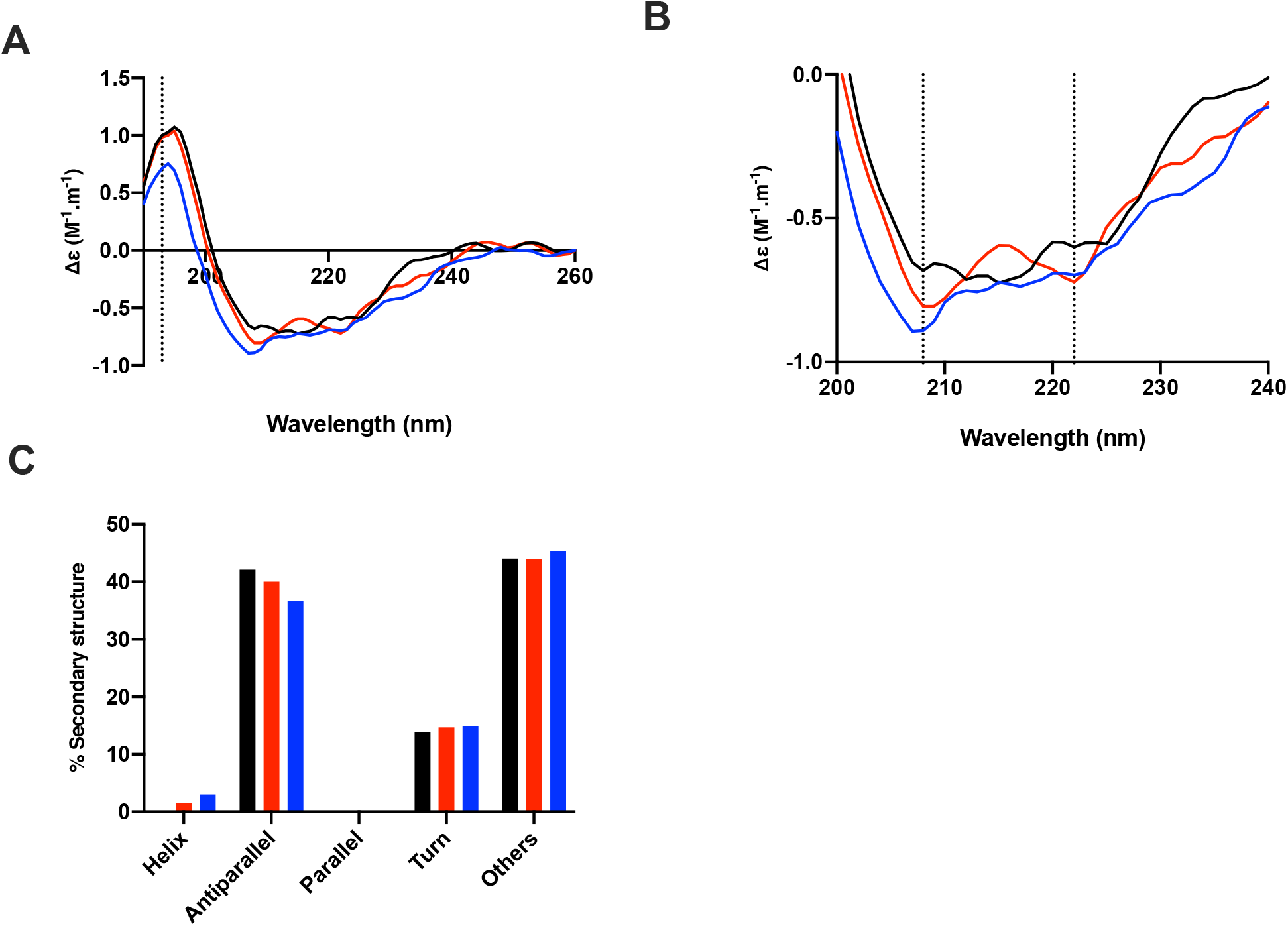
The conformational change of the SARS-CoV-2 S1 RBD observed in the presence of chondroitin sulphate D by circular dichroism (CD) spectroscopy. (**A**) Circular dichroism spectra (190 - 260 nm) of nCovS1RBD alone (black solid line), heparin (red solid line) and chondroitin sulphate D (blue) in PBS, pH 7.4. The dotted vertical line indicates 193 nm. (**B**) Details of the same spectra expanded between 200 and 240 nm. Vertical dotted lines indicate 222 nm and 208 nm. (**C**) Secondary structure content analysed using BeStSel for nCovS1RBD. α helical secondary structure is characterized by a positive band at ~193 nm and two negative bands at ~208 and ~222 nm (analysis using BeStSel was performed on smoothed data between 190 and 260 nm.

**Figure 6:**
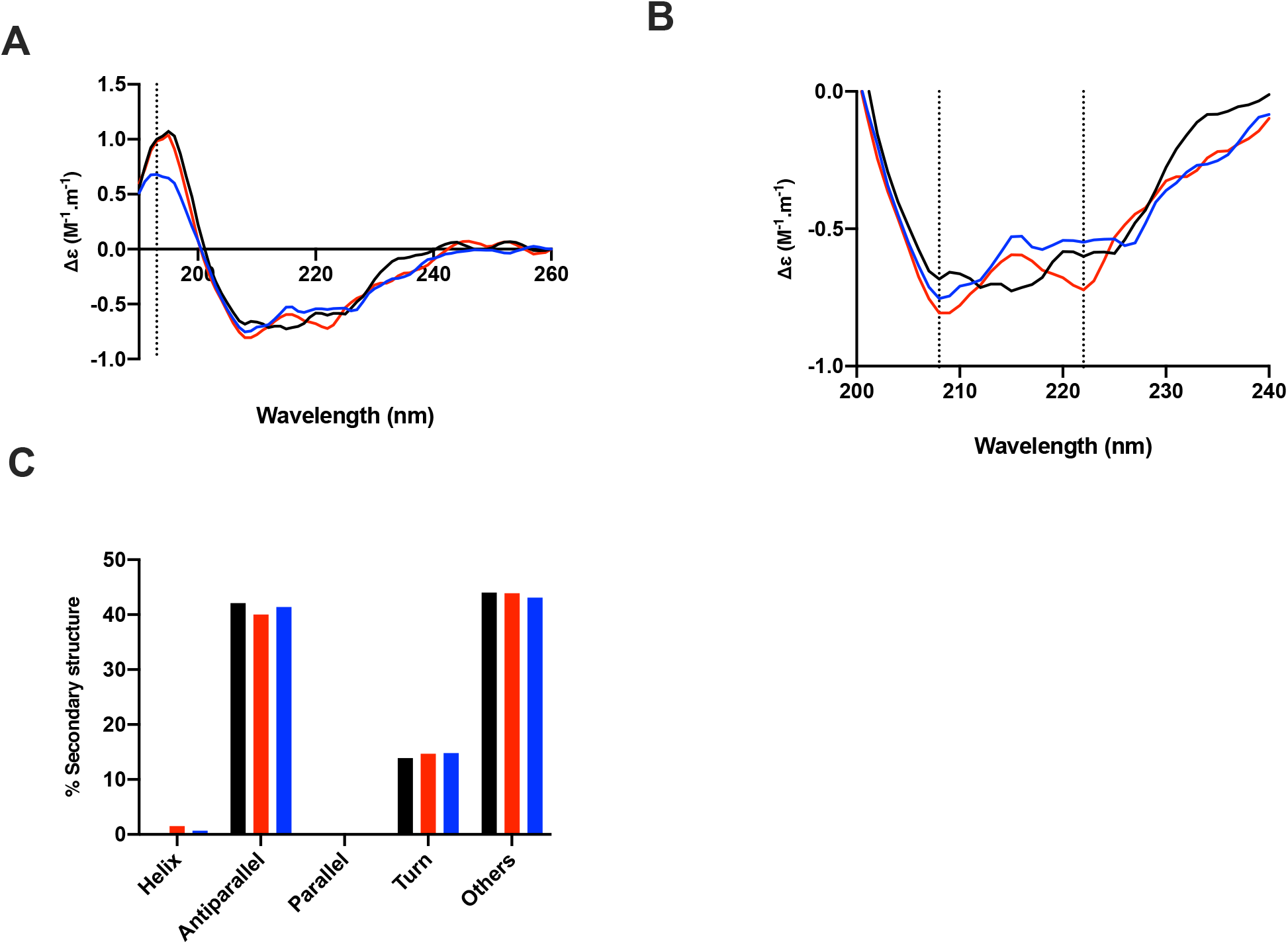
The conformational change of the SARS-CoV-2 S1 RBD observed in the presence of hyaluronic acid by circular dichroism (CD) spectroscopy. (**A**) Circular dichroism spectra (190 - 260 nm) of nCovS1RBD alone (black solid line), heparin (red solid line) and hyaluronic acid (blue) in PBS, pH 7.4. The dotted vertical line indicates 193 nm. (**B**) Details of the same spectra expanded between 200 and 240 nm. Vertical dotted lines indicate 222 nm and 208 nm. (**C**) Secondary structure content analysed using BeStSel for nCovS1RBD. α helical secondary structure is characterized by a positive band at ~193 nm and two negative bands at ~208 and ~222 nm (analysis using BeStSel was performed on smoothed data between 190 and 260 nm.

## Discussion and Conclusion

The rapid spread of SARS-CoV-2 represents a significant challenge to global health authorities. As there are currently no approved drugs to treat, prevent and/or mitigate the effects of SARS-CoV-2, repurposing existing pharmaceuticals and nutraceuticals is both a timely and appealing strategy. Glycosaminoglycans are generally well-tolerated and have been used successfully for many years with limited and manageable side effects.

Studying the structure and behaviour of SARS-CoV-2 Spike protein in solution is a vital step for the development of effective therapeutics against SARS-CoV-2. Here, the ability of the SARS-CoV-2 S1 RBD to interact with glycosaminoglycans has been studied using circular dichroism. The data show that SARS-CoV-2 S1 RBD interacts with GAGs other than UFH/LMWH and induces significant structural change.

Glycosaminoglycans are ubiquitously present on almost all mammalian cells and this class of carbohydrates are central to the strategy employed by *coronaviridae* to attach to host cells. Heparin has previously been shown to inhibit SARS-associated coronavirus cell invasion^2,3,15^ and this, in concert with the GAG-based data presented within this study, supports the utilisation of GAG-derived therapeutics against SARS-associated coronavirus.

It is noteworthy that the vast majority of commercially available GAG preparations remain a polydisperse mixture of natural products, containing both anticoagulant and non-anticoagulant saccharide structures. These may prove to be an invaluable resource for next-generation, biologically active, antiviral agents that display negligible anticoagulant potential, whilst the former remains tractable to facile, chemical (and enzymatic) engineering strategies to ablate their anticoagulation activities.

The further subfractionation of extant GAG preparations against anticoagulant activities (with proven low-toxicity profiles, good bioavailability and industrial-scale manufacturing) for off-label pathologies, provides an attractive strategy for quickly and effectively responding to COVID-19 and for the development of next generation GAG-based therapeutics.

Such drugs will be amenable to routine parenteral administration through currently established routes and additionally, direct to the respiratory tract via nasal administration, using nebulised heparin, which would be unlikely to gain significant access to the circulation. Thus, the anticoagulant activity of some GAGs, which can in any event be engineered out, would not pose a problem. Importantly, such a route of administration would not only be suitable for prophylaxis, but also for patients under mechanical ventilation^16^.

